# Modeling SARS-CoV-2 infection *in vitro* with a human intestine-on-chip device

**DOI:** 10.1101/2020.09.01.277780

**Authors:** Yaqiong Guo, Ronghua Luo, Yaqing Wang, Pengwei Deng, Min Zhang, Peng Wang, Xu Zhang, Kangli Cui, Tingting Tao, Zhongyu Li, Wenwen Chen, Yongtang Zheng, Jianhua Qin

## Abstract

Coronavirus disease 2019 (COVID-19) caused by severe acute respiratory syndrome coronavirus (SARS-CoV-2) has given rise to a global pandemic. The gastrointestinal symptoms of some COVID-19 patients are underestimated. There is an urgent need to develop physiologically relevant model that can accurately reflect human response to viral infection. Here, we report the creation of a biomimetic human intestine infection model on a chip system that allows to recapitulate the intestinal injury and immune response induced by SARS-CoV-2, for the first time. The microengineered intestine-on-chip device contains human intestinal epithelium (co-cultured human intestinal epithelial Caco-2 cells and mucin secreting HT-29 cells) lined in upper channel and vascular endothelium (human umbilical vein endothelial cells, HUVECs) in a parallel lower channel under fluidic flow condition, sandwiched by a porous PDMS membrane coated with extracellular matrix (ECM). At day 3 post-infection of SARS-CoV-2, the intestine epithelium showed high susceptibility to viral infection and obvious morphological changes with destruction of intestinal villus, dispersed distribution of mucus secreting cells and reduced expression of tight junction (E-cadherin), indicating the destruction of mucous layer and the integrity of intestinal barrier caused by virus. Moreover, the endothelium exhibited abnormal cell morphology with disrupted expression of adherent junction protein (VE-cadherin). Transcriptional analysis revealed the abnormal RNA and protein metabolism, as well as activated immune responses in both epithelial and endothelial cells after viral infection (e.g., up-regulated cytokine genes, TNF signaling and NF-kappa B signaling-related genes). This bioengineered *in vitro* model system can mirror the human relevant pathophysiology and response to viral infection at the organ level, which is not possible in existing *in vitro* culture systems. It may provide a promising tool to accelerate our understanding of COVID-19 and devising novel therapies.

## INTRODUCTION

COVID-19 caused by SARS-CoV-2 has become a global epidemic. As of late August 2020, it has caused 22 million infections and more than 790,000 deaths. SARS-CoV-2 infection is characterized by a process ranging from mild disease to severe systemic symptoms of multiple organs, most notably the lung, gastronintestinal tract, etc., and finally multi-organ failure (*1*). Respiratory symptoms dominate the clinical manifestations of COVID-19, yet obvious gastrointestinal symptoms were observed in 20% to 50% of patients, including abdominal pain, diarrhea, hematochezia and even intestinal perforation (*2-5*). In particular, these symptoms sometimes appear before the onset of respiratory symptoms (*6*). In addition, biopsy samples showed a large number of interstitial edema plasma cells and lymphocytes infiltrated into lamina propria of the stomach, duodenum and rectum (*7*). It has been reported that SARS-CoV-2 can cause acute hemorrhagic colitis, providing the evidence of the gastrointestinal tract implicating in the transmission of SARS-CoV-2 infection (*8*). Moreover, the viral RNA was identified in stool samples of COVID-19 patients and typical coronavirus virions in rectal tissue were observed under electron microscopy, implying that SARS-CoV-2 could potentially be transmitted via the fecal-oral route (*9-11*). These clinical evidences suggest that intestine represents another high-risk organ for SARS-CoV-2 infection besides the lungs, but the pathogenesis of intestinal infection underlying COVID-19 is elusive.

*In vivo*, human intestine contains complex multicellular components and host-pathogen interactions in a physiological flow microenvironment with mechanical cues. Currently, SARS-CoV-2 infection in intestine is studied based on monolayer cultures of intestinal epithelial cells (*12, 13*) and human organoids (*14, 15*). However, these *in vitro* models still have their limitations. The monolayer cell culture is oversimplified and cannot recapitulate the multiple cellular component, complex structure and functions of native intestine. Moreover, they lack cell-cell/matrix interactions and tissue-specific dynamic microenvironment that exist *in vivo*. Recently, intestinal organoids have provided a new *in vitro* 3D model for SARS-CoV-2 infection by providing multiple cell types and supporting viral replication in the gut enterocytes (*14*). But these organoids are still limited by the lack of typical characteristics of intestinal barrier, ECM, immune cells and physiological flow, which are key features of the intestinal microenvironment. As such, it is highly desirable to develop alternative *in vitro* models to better reflect the pathophysiology of human organs to SARS-CoV-2 infection.

Organ-on-a-chip technology has evolved to provide the possibility to reproduce the complex structures and physiological functions units of human organs in engineered microfluidic culture device (*16-18*). It has been utilized to represent the organ-level physiology and pathology, and applied in various biomedical applications, including organ engineering, disease studies and drug testing (*19-23*). In this work, we established a perfusable intestine infection model on chip system that simulate the human-relevant intestine response to SARS-CoV-2 infection at organ level. The microengineered intestine chip device consists of human intestine epithelium layer and vascular endothelium layer separated by an ECM coated porous PDMS membrane, in which Caco-2 cells and HT-29 cells are co-cultured in the upper channel, while HUVEC cells and circulating immune cells are lined in the lower channel under fluid flow. Using this system, we examined the infection and replication of SARS-CoV-2 in epithelial cells. We then systematically analyzed the changes of intestinal epithelium and endothelium induced by virus infection using confocal imaging. We also characterized the pathological changes and immune responses of intestinal barrier after viral infection via RNA-sequencing analysis. This human disease model on chip offers a novel strategy and platform for organ-level COVID-19 research and potential therapeutics.

## RESULTS

### Characterization of human intestine-on-chip device

*In vivo*, human intestinal epithelium is a cell monolayer that constitutes the intestinal barrier, which allows the mucus secretion, absorption of nutrients and prevents the invasion of pathogenic antigen or toxins in a dynamic microenvironment. In order to create an *in vitro* human model of intestine infection induced by SARS-CoV-2, we initially designed and constructed an intestine chip in a perfusable multilayer microfluidic culture device. Briefly, the device consists of two parallel channels separated by a thin and porous PDMS membrane coated with ECM to form bio-interface and facilitate the nutrient exchange between upper and lower cell layers.

In this work, the intestine epithelial cells Caco-2 and intestinal mucin secreting cells HT-29 were co-cultured to build intestinal epithelium in the upper channel under fluid flow (200 μL/h) to simulate the intraluminal fluid flow in intestine. The HUVEC cells were cultured in the lower channel to form vascular endothelium under fluid flow (50 μL/h). After 3 days of co-culture, the intestine epithelial and endothelial cells were confluent into monolayers under fluid flow, thus forming an intestine epithelium-endothelium tissue barrier. The integrity of the epithelial barrier was identified by assessing the expression of tight junction (ZO-1) and adherent junction (E-cadherin) proteins. The immunostaining analysis showed the positive expression of ZO-1 and E-cadherin in epithelium **(Fig. 2A)**. In particular, the epithelial cells formed villus-like structure lined by a highly polarized columnar epithelium identified by DIC imaging, which was similar to *in vivo* intestinal villi **(Fig. 2C)**. Moreover, the integrity of endothelium was examined by ZO-1 and VE-cadherin immunostaining, showing the regular and intact distribution of junction proteins **(Fig. 2B)**. As the intestinal mucus layer is the crucial interface between the host and microbes, which plays a vital role in preventing microbial invasion (*24*). We examined the mucus production in intestinal epithelium by immunostaining with mucin specific marker (MUC). As shown in **Fig. 2D**, it exhibited a scattered distribution of MUC positive cells in the intestinal epithelium, suggesting the secretory function of mucus. It appears that the established intestine chip system exhibited intestinal villus-like structure, the integrity of tissue barrier and ability of mucus secretion under fluidic flow, representing the key features of human intestinal barrier in a physiologically relevant manner.

**Figure 1.**
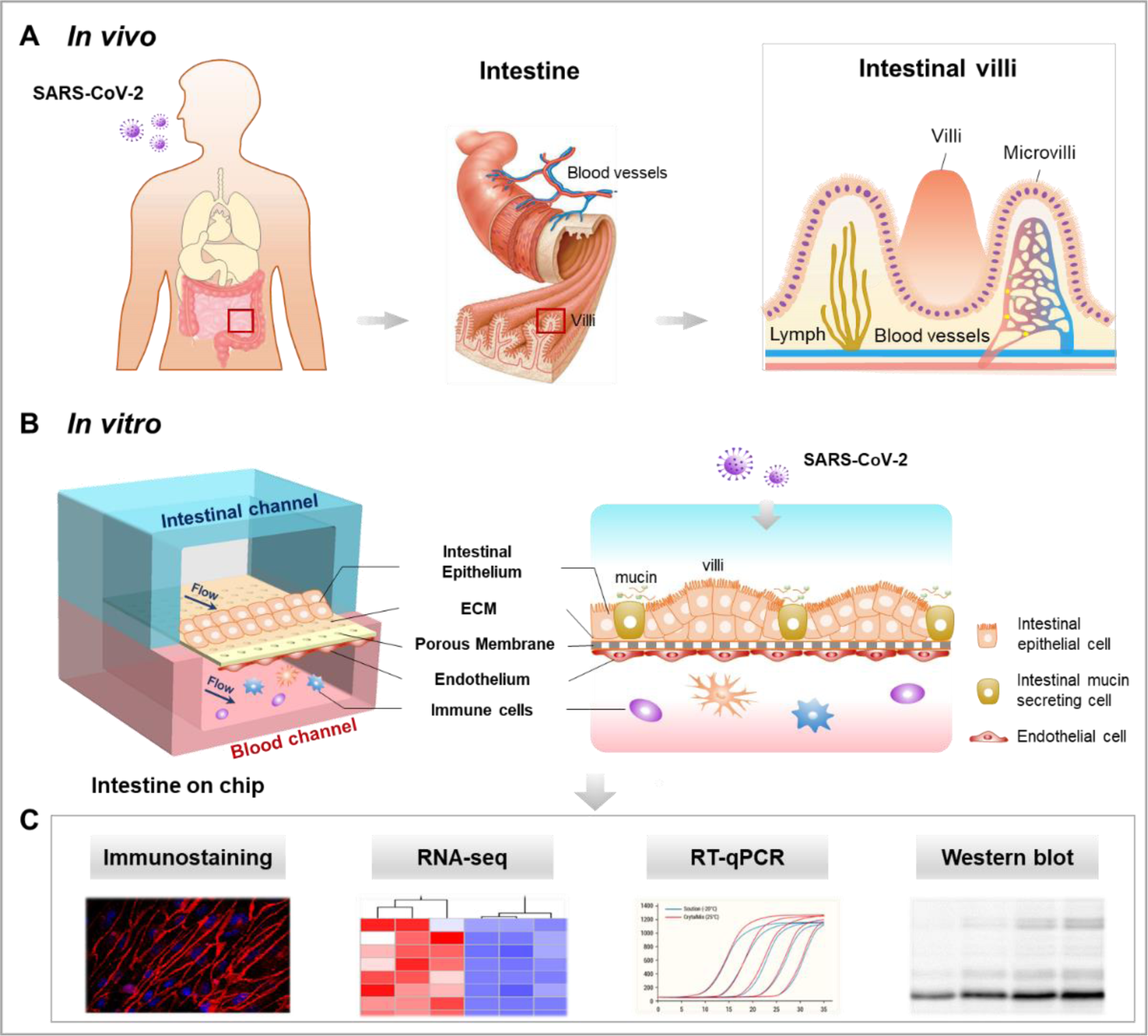
Schematic diagram of SARS-CoV-2-induced intestinal infection model on chip. **(A)** Illustration of SARS-CoV-2 infection of human small intestine *in vivo*. **(B)** The configuration SARS-CoV-2 induced intestinal infection model on chip. The device consists of upper intestinal epithelial channel (blue) and lower microvascular endothelial channel (red) separated by an ECM coated porous PDMS membrane. The intestinal barrier was established by co-culture of intestinal epithelial Caco-2 cells and intestinal mucin secreting HT-29 cells on the top channel. Human peripheral blood mononuclear cells (PBMC) were introduced to the bottom vascular channel during the progression of virus infection. **(C)** The responses of the intestinal chip to SARS-CoV-2 infection are analyzed using various methods.

**Figure 2.**
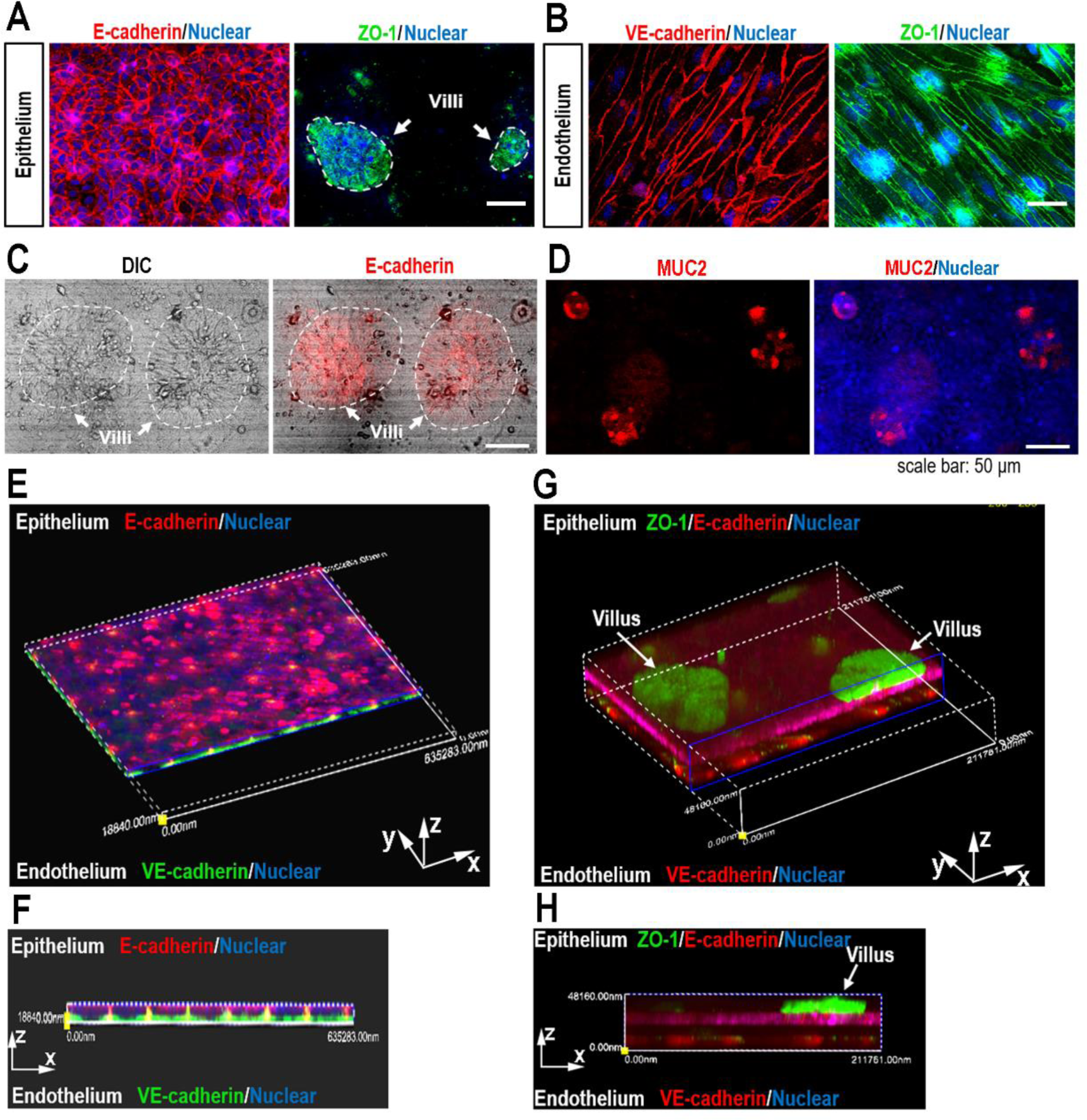
Characterization of intestinal epithelium and endothelium in the microengineered intestine chip. **(A)** Confocal micrographs showed the epithelial adhesion junctions visualized by E-cadherin and tight junctions visualized by ZO-1 in the intestine model. The intestinal villus-like structures with higher expression of ZO-1 were indicated by white dotted lines. **(B)** Confocal micrographs showed the endothelial adhesion junctions visualized by VE-cadherin and tight junctions visualized by ZO-1 in the intestine model. **(C)** The DIC image showed the intestinal villus-like structure with clumps of cells (indicated by white dotted line). **(D)** Confocal micrographs showed scattered MUC-positive cells on endothelium. Scale bar: 50 μm. **(E)** 3D reconstructed confocal image showed intestinal epithelium (E-cadherin) and endothelium (VE-cadherin) in the intestine model. **(F)** Side view of intestinal epithelium (E-cadherin) and endothelium (VE-cadherin) in the intestine model. **(G)** 3D reconstructed confocal image showed intestinal epithelium (E-cadherin), endothelium (VE-cadherin) and intestinal villus-like structure (indicated by white arrow) in the microengineered intestine model. **(H)** Side view of intestinal epithelium (E-cadherin), endothelium (VE-cadherin) and intestinal villus-like structure (indicated by white arrow) in the microengineered intestine model. Each image represents 3 independent experiments.

### SARS-CoV-2 infection in the intestine-on-chip

Prior to SARS-CoV-2 infection on the intestine chip, we first detected the expression level of ACE2 and TMPRSS2 in three cell lines by western blot, including Caco-2, HT-29 and HUVEC cells. It is recognized that SARS-CoV-2 uses ACE2 as a receptor for cellular entry and TMPRSS2 protease for viral Spike protein priming (*25, 26*). The intestinal enterocytes have a higher expression of ACE2 compared to alveolar epithelial type II cells (*27-29*). Among the tested three cell lines in this work, ACE2 and TMPRSS2 were expressed at the highest level in Caco-2 cells **(Fig. S1A)**, which implied that Caco-2 cells may be more susceptible to SARS-CoV-2 infection. Caco-2 cells were then inoculated with different multiplicity of virus (MOI=0.04, 0.4 and 2) to identify the virus susceptibility. Three days after infection, immunostaining data showed more than 80% Spike protein-positive cells were observed in Caco-2 cells at MOI of 2 **(Fig. S1B)**.

To examine the SARS-CoV-2 infection on the intestine chip, the virus was inoculated into the intestinal channel at MOI=1 for 24h after co-culture of intestinal epithelium and endothelium for 3 days. The virus infection in the intestinal epithelium layer and endothelium layer were then examined by immunostaining analysis. The data showed that Spike positive cells were predominantly detected on the intestinal layer, indicating that the intestinal epithelial cells are more susceptible to SARS-CoV-2 infection than endothelial cells **(Fig. 3)**.

**Figure 3.**
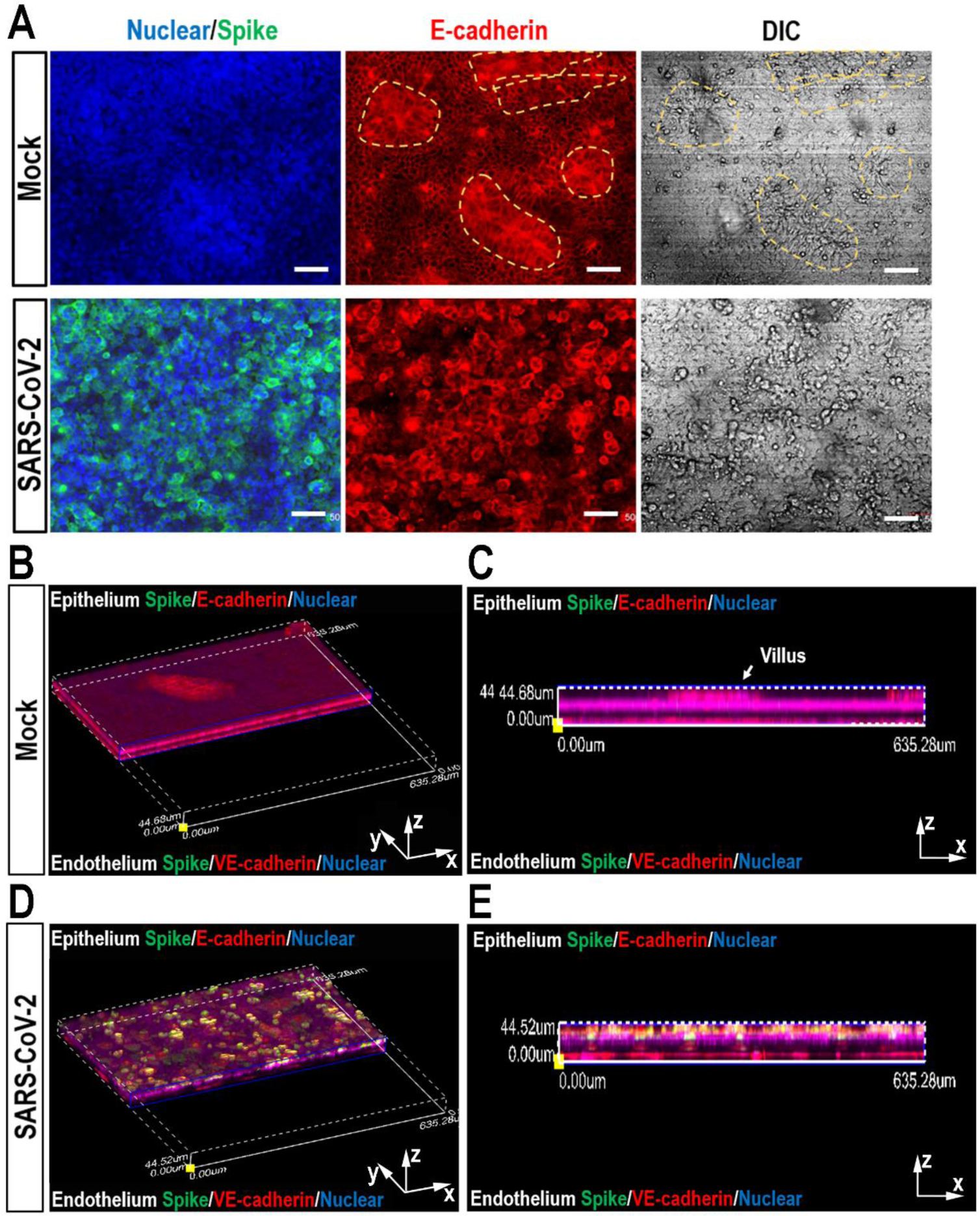
Examination of SARS-CoV-2 infection on the human intestine chip. **(A)** Confocal micrographs showed the effects of SARS-CoV-2 infection (Spike protein) on the intestinal epithelium (E-cadherin) and intestinal villus-like structure (indicated by yellow dotted line) of chip at day 3 post-infection following introduction of PBMCs into microvascular channel. **(B)** 3D reconstructed confocal image of mock-infected intestine model following introduction of PBMCs into microvascular channel. **(C)** Side view of the mock-infected intestine model. **(D)** 3D reconstructed confocal image of SARS-CoV-2-infected intestine model following introduction of PBMCs into microvascular channel. **(E)** Side view of the SARS-CoV-2-infected intestine model following introduction of PBMCs into microvascular channel. SARS-CoV-2 infection was predominantly identified in epithelial layer by viral Spike protein expression. Each image represents 3 independent experiments.

*In vivo*, the development and barrier maintenance of intestine depends heavily on the proper function of adherent junctions, we further examined the adherent junctions of intestinal epithelium and endothelium, respectively. Confocal micrographs of intestinal epithelium showed that on day 3 after viral infection, the adherent junction identified by E-cadherin was severely destroyed, accompanied by the damage of intestinal villus-like structures **(Fig. 3A)**. Moreover, the mucous secreting cells in intestinal epithelium were examined by immunostaining with MUC2. The results showed that the clonal distribution MUC2^+^ cells became dispersed after viral infection **(Fig. 4A)**. This suggests that the disturbed intestinal mucus layer was caused by viral infection, which may lead to virus encroachment and infection. It is noted that, in the vascular side, the adherent junctions between endothelial cells were severely disrupted by VE-cadherin immunostaining **(Fig. 4B)**. Quantitative analysis showed that the density and size of endothelial cells were significantly decreased **(Fig. 4C and 4D)**. It reveals the injury of vascular endothelial cells following viral infection, while there were no obvious Spike positive cells on the vascular side. The presence of endothelial injury might partially explain the pathogenesis of COVID-19 associated coagulopathy or vascular thrombosis (*30*).

**Figure 4.**
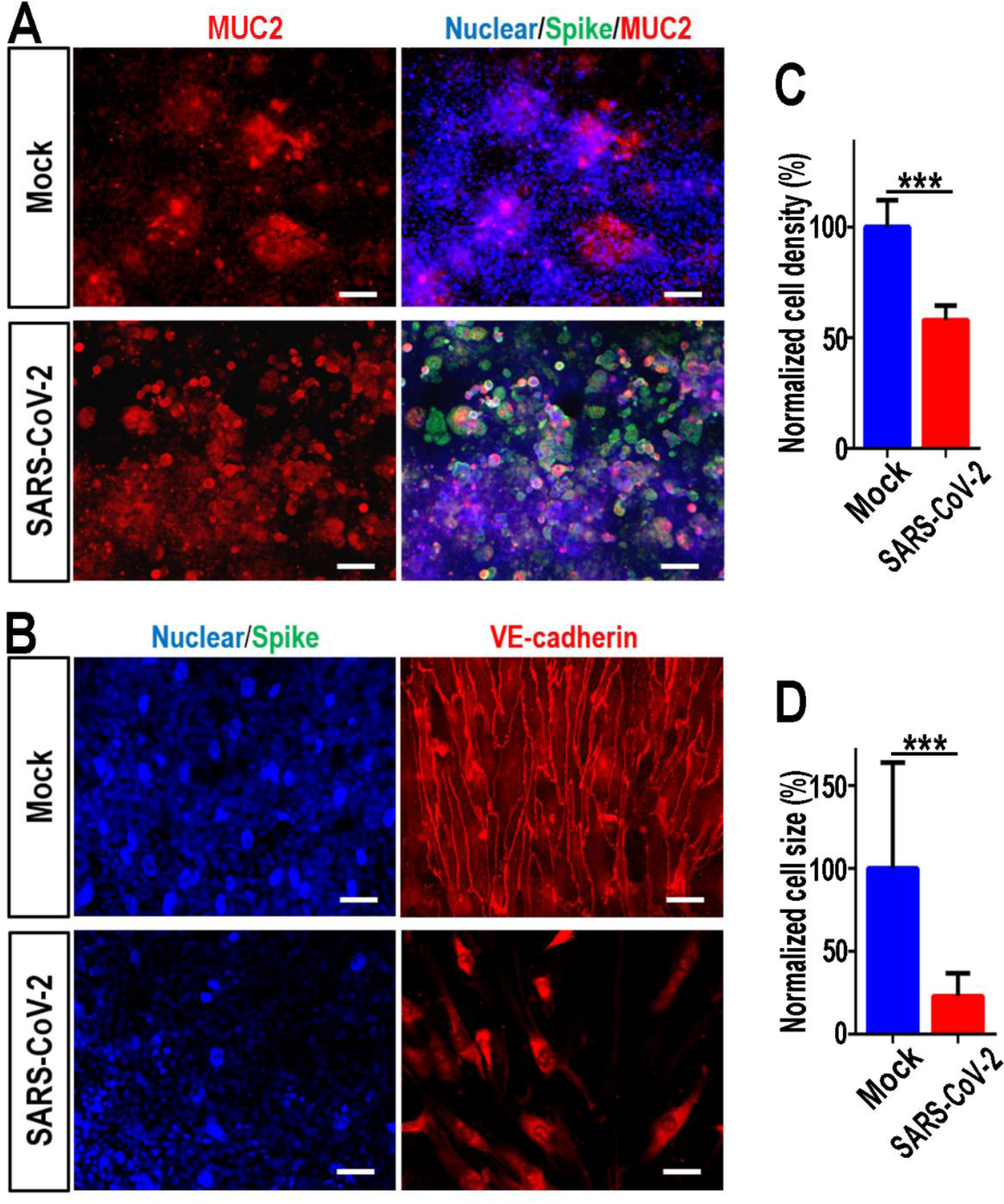
Morphological changes of intestinal barrier after exposure to SARS-CoV-2. **(A)** Confocal micrographs showed the effects of SARS-CoV-2 infection (Spike protein) on the epithelial MUC expression in the intestine model at day 3 post-infection following introduction of PBMCs into microvascular channel. **(B)** Confocal micrographs showed the effects of SARS-CoV-2 infection (Spike protein) on the intestinal endothelium (E-cadherin) in microengineered intestine model following introduction of PBMCs into microvascular channel. **(C, D)** Quantification of endothelial cell density and size for mock- and SARS-CoV-2-infected chips. 4 chips were counted for cell density quantification in each group, and 100 cells were counted for cell size quantification in each group. Data were presented as mean ± SD. Data were analyzed by Student’s t-test (***: p<0.001).

### Transcriptional analysis of host cells to SARS-CoV-2 infection

To gain a comprehensive overview of transcriptional responses to SARS-CoV-2 infection, we performed RNA-sequencing analysis of intestinal epithelial cells and endothelial cells in human intestine model. Briefly, 3 days after infection, intestinal epithelial cells and endothelial cells were collected separately and analyzed by RNA-sequencing. Volcano plots showed that SARS-CoV-2 infection induced dramatic transcriptome modulations in both intestinal epithelial cells and endothelial cells **(Fig. 5A)**. In order to identify differentially expressed genes (DEGs), the critical value and P value of the abundance fold change were set to 2.0 and 0.05, respectively. Among the DEGs, 9684 genes (4022 down-regulated genes and 5662 up-regulated genes) were significantly modulated in intestinal epithelial cells, while 6713 genes (2569 down-regulated genes and 4144 up-regulated genes) were significantly modulated in endothelial cells following viral infection. This result suggested that SARS-CoV-2 infection had a greater effect on intestinal epithelial cells than endothelial cells, possibly due to higher viral load in intestinal epithelium. By combining the two data sets, we found that intestinal epithelial cells and endothelial cells shared 1791 overlapping up-regulated DEGs (37.3% of total up-regulated DEGs) and 3317 overlapping down-regulated DEGs (51.5% of total down-regulated DEGs) **(Fig. 5B)**.

**Figure 5.**
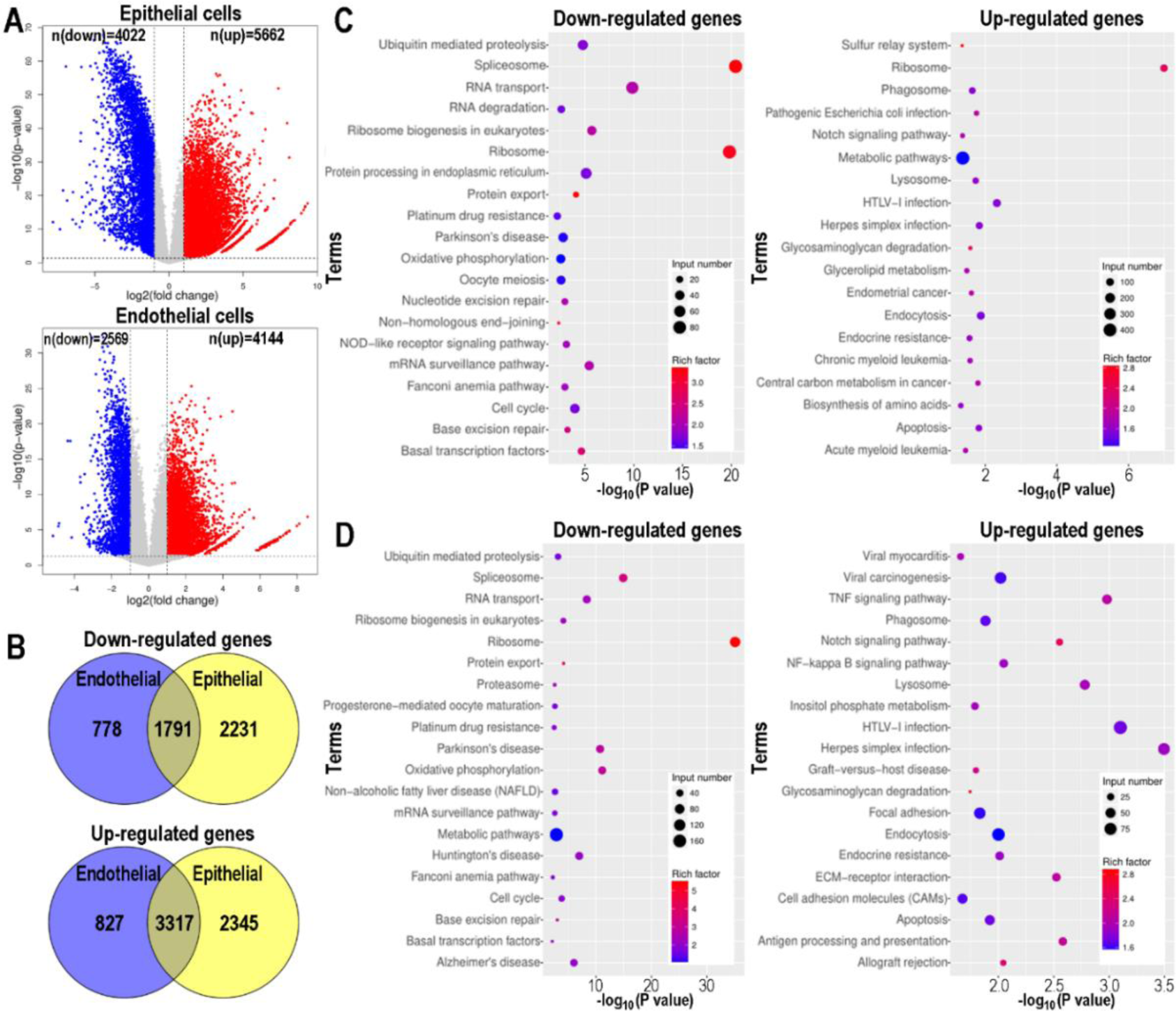
Transcriptional analysis of intestinal epithelial cells and endothelial cells to SARS-CoV-2 infection on chip. **(A)** Volcano plots showed the regulated genes of cells following SARS-CoV-2 infection. Genes differentially expressed with fold change over 2.0 and p<0.05 were marked in color. P values were calculated using a two-sided, unpaired student’s t-test with equal variance assumed (n=3 independent biological samples). **(B)** Venn diagrams depicting the differentially expressed genes shared or unique between each comparison. **(C)** KEGG pathway enrichment analysis of differentially expressed genes in intestinal epithelial cells following SARS-CoV-2 infection. **(D)** KEGG pathway enrichment analysis of differentially expressed genes in endothelial cells following SARS-CoV-2 infection. **(E, F)** The color of the dots represents the rich factor and the size represents the input number for each KEGG term. The horizontal axis indicates the significance of enrichment (-log10 (P value)), and the vertical axis indicates the enriched KEGG pathway (20 most enriched terms).

To investigate the host biological pathways modified by SARS-CoV-2 infection, KEGG pathway enrichments analysis comparing mock-infected versus SARS-CoV-2 infected intestine chips was performed **(Fig. 5C and 5D)**. The results revealed that the viral infection resulted in the abnormal pathway networks in both cell types, including RNA metabolism pathways (e.g., spliceosome, RNA transport, RNA degradation, mRNA surveillance pathway) and protein metabolism pathways (e.g., ubiquitin mediated proteolysis, protein export). Given the vital roles of RNA and protein metabolism in maintaining the normal physiological functions of cells, we reasoned that SARS-CoV-2 may seriously affect the host cells by disrupting these critical pathways. In addition, we found that some immune responses related pathways, such as TNF signaling pathway and NF-kappa B signaling pathway, were particularly enriched in genes that are significantly up-regulated in endothelial cells, which indicated that the intestinal endothelial cells play critical roles in mediating intestinal immune responses.

### Immune response of intestinal model on chip to SARS-CoV-2 infection

According to the above transcriptional analysis, the KEGG pathway enrichment analysis indicated some immune response-related signaling pathways were activated following SARS-CoV-2 infection. We then tried to identify the up-regulated cytokine genes that mediated the subsequent inflammatory responses underlying SARS-CoV-2-induced intestinal infection. The heat map showed that SARS-CoV-2 infection elicited an extensive cytokines induction in both intestinal epithelium and endothelium, including TNF, interleukins, chemokines and colony-stimulating factors in both intestinal epithelial cells and endothelial cells **(Fig. 6A and 6B)**. Notably, in this infected intestinal model, some cytokines including TNF, IL-6, CXCL10, CCL5 and CSF3 were significantly up-regulated, which is similar to the clinical manifestations of patients with severe COVID-19 (*31, 32*). These results may partially explain the presence of infiltrated immune cells and inflammatory response in the intestine of patients with severe COVID-19.

**Figure 6.**
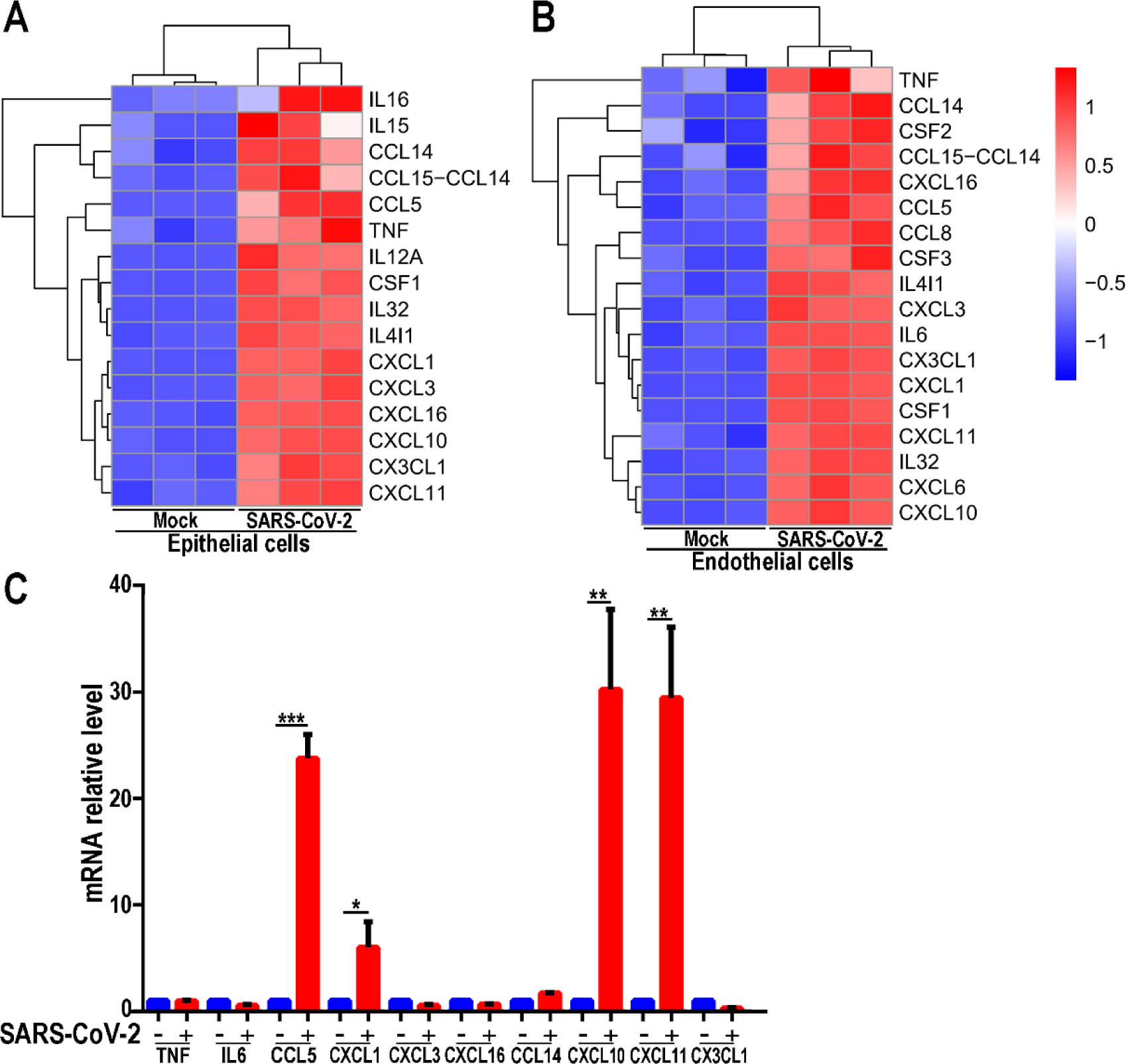
Assessment of immune responses in human intestine chip following SARS-CoV-2 infection. **(A)** Heatmaps depicting the significantly up-regulated cytokine genes related in intestinal epithelial cells. **(B)** Heatmaps depicting the significantly up-regulated cytokine genes related in endothelial cells. Colored bars represent Z-score of log2 transformed values. **(C)** The relative mRNA level of indicated genes were examined by qRT-PCR. Data were presented as mean ± SD. Data were analyzed by Student’s t-test (*: p<0.05; **: p<0.01; ***: p<0.001). N=3.

Then, we further detected the expression of pro-inflammatory cytokines and chemokines-related genes in the intestinal epithelial cells by qRT-PCR. The expression of CCL5, CXCL1, CXCL10 and CXCL11 were significantly up-regulated in the SARS-CoV-2-infected epithelial cells **(Fig. 6C)**. The data indicated that chemokine genes could be induced by SARS-CoV-2, which may play a crucial role in the recruitment of immune cells and modulation of immune response to virus infection. It has been previously reported that chemokines CCL5 can recruit T cells, dendritic cells, eosinophils, NK cells, mast cells, and basophils (*33*), CXCL1 recruits neutrophils (*34*), CXCL10 recruits activated Th1 lymphocytes (*35*), and CXCL11 recruits interleukin-activated T-cells (*36*) to sites of inflammation. This might provide the clues to further identify which immune cells are involved in the intestinal inflammatory responses in the progression of COVID-19.

## DISCUSSION

In this study, we created a biomimetic human disease model of intestinal infection on chip by SARS-CoV-2, which could mirror the pathophysiological features and immune response associated with gastrointestinal symptoms in COVID-19. This *in vitro* model can reflect the damage of the intestinal epithelial barrier, including the destruction of intestinal villi, the disruption of barrier integrity, mucosal secretion disorder and activated immune responses after SARS-CoV-2 infection, which may help us to understand the pathological processes involved in COVID-19. Moreover, despite the low susceptibility to virus infection, the vascular endothelium displayed significant morphological damage induced by SARS-CoV-2, revealing the possible complex cross-talk between intestine epithelial cells and endothelial cells in mediating tissue barrier injury and COVID-19 progression.

*In vivo*, human intestine comprises the largest component of human immune system with large populations of scattered innate and adaptive effector cells, because it is constantly exposed to foreign antigens and other environmental agents. The protective mucus gel secreted by goblet cells is distributed on the gastrointestinal epithelium, which provides chemical and physical defenses for the intestinal barrier and plays a crucial role in intestinal homeostasis. In this intestine chip, the Caco-2 cells and mucin secreting HT-29 cells were co-cultured under continuous perfusion to form intestinal epithelium, which can recapitulate the intestinal barrier in a highly physiological relevant manner. In particular, we found that the secretion of intestinal mucin has undergone unrecognized changes, from agglomerated distribution to scattered distribution after viral infection. It is speculated that the disturbed mucus layer may be associated with further invasion and infection of virus. These results may also explain the increased intestinal permeability and further cause symptoms, such as diarrhea^6^ and hemorrhagic colitis (*37*) in COVID-19 patients.

Transcriptome analysis demonstrated significantly altered vital biological processes in both intestinal epithelium and endothelium following SARS-CoV-2 infection, including RNA and protein metabolism pathways, cell cycle regulation and oxidative phosphorylation. Specifically, we identified many up-regulated cytokine and chemokine-related genes in the intestinal epithelial cells, similar to the clinical manifestations of COVID-19. Because chemokine can act as chemoattractant to recruit immune cells to the infected sites, these altered chemokines may contribute to the immune cells mediated inflammatory responses in the intestine. In addition, we found that the intestinal epithelium is more susceptible to SARS-CoV-2 infection than endothelium on chip, which is consistent with previous studies (*12*). As such, we assume that the vascular endothelial injury may be mediated by the inflammatory factors or paracrine signals released by virus infected intestinal epithelial cells.

One potential limitation of this work is the use of immortalized intestinal epithelial cell lines (e.g., Caco-2) originally isolated from human colon tumors. However, Caco-2 cells have been widely used to recapitulate many physiological and pathological functions of human intestine *in vivo (38, 39)*. The intestinal epithelium differentiated from primary intestinal stem cells may be selected for further studies in later time.

Overall, in this work, we provide the proof-of-concept to build an intestine-on-chip infection model at organ-level that permits to closely mirror the intestinal pathophysiology and human response to SARS-CoV-2 infection, which is impossible to achieve by existing *in vitro* culture models. It can not only supply a unique, rapid and low-cost *in vitro* platform for viral infection, but also provide a complement to animal models to study disease progress, virus transmission and host-virus interactions in a more realistic manner, thereby accelerating the COVID-19 research and development of novel therapeutics.

## MATERIALS AND METHODS

### Device fabrication

The human intestine model consisted of the upper and lower layers fabricated by using conventional soft lithography procedures. Polydimethylsiloxane (PDMS) pre-polymer was prepared by mixing 10:1 (wt/wt) PDMS base to curing agent (184 Silicone Elastomer, Dow Corning Corp) and casted on molds to produce molded device with channels by thermal curing at 80 °C for 45 min. The channels were 1.5 mm wide × 0.25 mm high and the length of overlapping channels was 8 mm. The two channels were separated by a thin (∼20 μm) through-hole PDMS membrane (pore size: 5 μm) to construct tissue-tissue interfaces. The porous PDMS membranes were fabricated based on the glass templates and spin-coating method modified from the previous protocol. The membrane was sandwiched in the intestine-on-chip device by oxygen plasma bonding. Finally, the devices were exposed to ultraviolet light for disinfection and sterilization. Prior to cell seeding, both sides of the porous membrane were coated with rat tail collagen type I (200 μg/mL, Corning) and incubated at 37 °C for 48 h.

### Cell culture

Human colon adenocarcinoma cells (Caco-2) were cultured in high-glucose Dulbecco’s Modified Eagle’s Medium (DMEM, Gibco) supplemented with 10% Fetal Bovine Serum (FBS, Gibco). Human colorectal adenocarcinoma grade II cells HT29 were purchased from Procell Life Science&Technology Co., Ltd (Procell, CL-0118) and were maintained in growth medium (Procell, CM-0118) supplemented with 10% FBS. Human umbilical vein endothelial cells (HUVEC) were isolated from human umbilical cord and cultured in Endothelial Cell Medium purchased from ScienCell Research Laboratories, inc. Human peripheral blood mononuclear cells were isolated from fresh human blood using Ficoll (Stem cell technologies) density centrifugation. Isolated PBMCs were resuspended in RPMI 1640 medium containing 10% FBS and 50 IU/mL IL-2 and used for adhesion assays on chip. All cells were cultured at 37 °C in a humidified atmosphere of5% CO2.To create the intestine model, HUVEC cells (∼ 1×10^5^ cells) were initially seeded on the bottom side of the collagen-coated porous PDMS membrane and allowed to attach on the membrane surface under static conditions for two hours. Subsequently, cells were washed with fresh medium to exclude/remove unattached ones. Then, Caco-2 cells (∼ 1×10^5^ cells) and HT29 cells (∼ 1×10^4^ cells) were mixed and seeded into the upper channel under static cultures. After cell attachment, cells were grown to confluence for 3 days and the chips were maintained in an incubator with 5% CO2 at 37°C.

### Virus

A clinical isolate SARS-COV-2 strain 107 was obtained from Guangdong Provincial Center for Disease Control and Prevention, China, and propagated in Vero E6 cells. The virus titers were (infectious titers of virus) were determined by a TCID50 assay on Vero cells. All work involving live SARS-CoV-2 was performed in the Chinese Center for Disease Control and Prevention-approved BSL-3 laboratory of the Kunming Institute of Zoology, Chinese Academy of Science.

### SARS-CoV-2 infection

Caco-2 cells were seeded in 24-well plates (2×10^5^ cells per well) in high-glucose Dulbecco’s Modified Eagle’s Medium containing 10% FBS. After seeding for 24h, cells were infected with SARS-CoV-2 at an indicated multiplicity of virus (MOI=0.04, 0.4 and 2). After one hour, cells were washed three times with PBS and kept in fresh medium for 3 days. At day 3 post-infection, cells were washed with PBS and then fixed with 4% paraformaldehyde (PFA) before analysis.

For SARS-CoV-2 infection in the human intestine model, the apical channel of chip device was infused with 30 μL of high-glucose DMEM medium containing the indicated multiplicity of virus (MOI=1). After one hour of infection, cells were washed three times with PBS and kept in fresh medium. At day 3 post-infection, the epithelial cells and endothelial cells cultured on chip were fixed for immunofluorescence analysis or lysed for RNA-sequencing analysis, respectively.

### Immunostaining

Caco-2 cells cultured on well plate were washed with PBS and fixed with 4% PFA at 4°C overnight. Cells were then permeabilized with 0.2% Triton X-100 in PBS (PBST buffer) for 5 min and blocked with PBST buffer containing 5% normal goat serum for 30 minutes at room temperature. Antibodies were diluted with PBST buffer. Cells were stained with corresponding primary antibodies at 4°C overnight and with secondary antibodies (supplementary Table S1) at room temperature for 1hour. After staining with secondary antibodies, cell nuclei were counterstained with DAPI. For immunofluorescent imaging of intestine model, cells were washed with PBS through the upper and bottom channels and fixed with 4% PFA. The fixed tissues were subjected to immunofluorescence staining by the same procedure as described above. All images were acquired using a confocal fluorescent microscope system (FV-1000, Olympus). Image processing was done using ImageJ (NIH).

### RNA extraction, library preparation and sequencing

Intestinal epithelial cells and endothelial cells were collected separately from the chips, and total RNAs were extracted from samples using Trizol (Invitrogen). DNA digestion was carried out after RNA extraction by DNaseI. RNA quality was determined by examining A260/A280 with NanodropTM OneCspectrophotometer (Thermo Fisher Scientific Inc). RNA Integrity was confirmed by 1.5% agarose gel electrophoresis. Qualified RNAs were finally quantified by Qubit3.0 with QubitTM RNA Broad Range Assay kit (Life Technologies). 500 ng total RNAs were used for stranded RNA sequencing library preparation using KC-DigitalTM Stranded mRNA Library Prep Kit for Illumina® (Catalog NO. DR08502, Wuhan SeqHealth Co., Ltd. China) following the manufacturer’s instruction. The kit eliminates duplication bias in PCR and sequencing steps, by using unique molecular identifier (UMI) of 8 random bases to label the pre-amplified cDNA molecules. The library products corresponding to 200-500 bps were enriched, quantified and finally sequenced on Hiseq X 10 sequencer (Illumina).

### RNA-sequencing analysis

Raw sequencing data was first filtered by Trimmomatic (version 0.36), low-quality reads were discarded and the reads contaminated with adaptor sequences were trimmed. Clean Reads were further treated with in-house scripts to eliminate duplication bias introduced in library preparation and sequencing. In brief, clean reads were first clustered according to the UMI sequences, in which reads with the same UMI sequence were grouped into the same cluster, resulting in 65,536 clusters. Reads in the same cluster were compared to each other by pairwise alignment, and then reads with sequence identity over 95% were extracted to a new sub-cluster. After all sub-clusters were generated, multiple sequence alignment was performed to get one consensus sequence for each sub-cluster. After these steps, any errors and biases introduced by PCR amplification or sequencing were eliminated.

The de-duplicated consensus sequences were used for standard RNA-sequencing analysis. They were mapped to the reference genome of *Homo sapiens* from Ensembl database (ftp://ftp.ensembl.org/pub/release-87/fasta/homo_sapiens/dna/) using STAR software (version 2.5.3a) with default parameters. Reads mapped to the exon regions of each gene were counted by featureCounts (Subread-1.5.1; Bioconductor) and then RPKMs were calculated. Genes differentially expressed between groups were identified using the edgeR package (version 3.12.1). An FDR corrected p-value?cutoff of 0.05 and Fold-change cutoff of 2 were used to judge the statistical significance of gene expression differences. Gene ontology (GO) analysis and Kyoto encyclopedia of genes and genomes (KEGG) enrichment analysis for differentially expressed genes were both implemented by KOBAS software (version: 2.1.1) with a corrected P-value cutoff of 0.05 to judge statistically significant enrichment.

### Statistical analyses

Data were collected in Excel (Microsoft). Differences between two groups were analyzed using a Student’s t-test. Multiple group comparisons were performed using a one-way analysis of variance (ANOVA) followed by post-hoc tests. The bar graphs with error bars represent mean ± standard deviation (SD). Significance is indicated by asterisks: *, p < 0.05; **, p < 0.01; ***, p < 0.001.

## Data availability

All relevant data are available in the manuscript or supporting information. All of the RNA-sequencing raw data have been deposited on SRA under the accession number PRJNA658711.

## Supporting information

Supporting information is available from the Wiley Online Library or from the author.

## Acknowledgments

This research was supported by the Strategic Priority Research Program of the Chinese Academy of Sciences, Grant (Nos. XDB29050301, XDA16020900, XDB32030200), National Key R&D Program of China (No. 2017YFB0405404), The National key Research and Development Program of China (2020YFC0842000), National Science and Technology Major Project (No. 2018ZX09201017-001-001), National Nature Science Foundation of China (Nos. 31671038, 31971373, 81703470, 81803492), China Postdoctoral Science Foundation (No. 2019M660065), Innovation Program of Science and Research from the DICP, CAS (DICP I201934). We thank Prof. Yonggang Yao (Kunming Institute of Zoology, CAS) for his strong support on this work.

