## Supplementary Figure 1 for "Modeling SARS-CoV-2 infection *in vitro* with a human intestine-on-chip device"

**Supplementary materials**

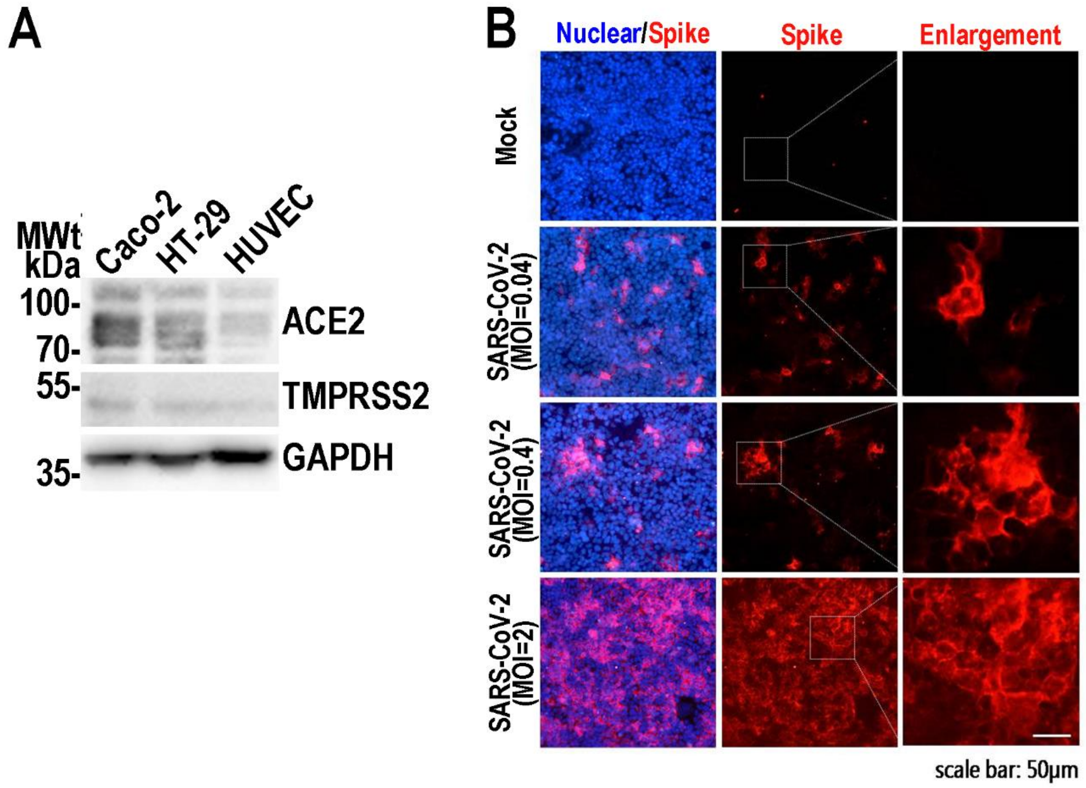


**Supplementary Figure 1. Examination of SARS-CoV-2 infection in Caco-2 cells in monolayer culture. (A)** Western blot was perform to detect the expression level of ACE2 for 4 cell lines. **(B)** Immunofluorescent images showed the viral Spike protein (Spike protein S1 subunit of SARS-CoV-2) staining in Caco-2 cells at 72-hour post SARS-CoV-2 infection at an indicated multiplicity of virus (MOI=0.04, 0.4 and 2).
